# Experimental increases in glucocorticoids alter function of the HPA axis in wild red squirrels without negatively impacting survival and reproduction

**DOI:** 10.1101/309278

**Authors:** Freya van Kesteren, Brendan Delehanty, Sarah E. Westrick, Rupert Palme, Rudy Boonstra, Jeffrey E. Lane, Stan Boutin, Andrew G. McAdam, Ben Dantzer

**Affiliations:** Department of Psychology, University of Michigan, MI 48109 Ann Arbor, Michigan, USA; Department of Biological Sciences, University of Toronto Scarborough, Toronto, Ontario, M1C 1A4 Canada; Department of Biomedical Sciences, University of Veterinary Medicine, A-1210 Vienna, Austria; Department of Biology, University of Saskatchewan, Saskatoon, SK S7N 5E2, Canada; Department of Biological Sciences, University of Alberta, Edmonton, Alberta, Canada T6G 2E9; Department of Integrative Biology, University of Guelph, Guelph, Ontario, Canada N1G 2W1; Department of Ecology and Evolutionary Biology, University of Michigan, MI 48109, Ann Arbor, Michigan, USA

**Keywords:** cortisol, glucocorticoids, hormone manipulations, hypothalamic–pituitary–adrenal (HPA) axis, North American red squirrels, stress

## Abstract

Hormones such as glucocorticoids (colloquially referred to as “stress hormones”) have important effects on animal behavior and life history traits, yet most of this understanding has come through correlative studies. While experimental studies offer the ability to assign causality, there are important methodological concerns that are often not considered when manipulating hormones, including glucocorticoids, in wild animals. In this study, we examined how experimental elevations of cortisol concentrations in wild North American red squirrels (*Tamiasciurus hudsonicus*) affected their hypothalamic–pituitary–adrenal (HPA) axis reactivity, and life history traits including body mass, litter survival, and adult survival. The effects of exogenous cortisol on plasma cortisol concentrations depended on the time between treatment consumption and blood sampling. In the first nine hours after consumption of exogenous cortisol, individuals had significantly higher true baseline plasma cortisol concentrations, but adrenal gland function was impaired as indicated by their dampened response to capture and handling and to injections of adrenocorticotropic hormone compared to controls. Approximately 24 hours after consumption of exogenous cortisol, individuals had much lower plasma cortisol concentrations than controls, but adrenal function was restored. Corticosteroid binding globulin (CBG) concentrations were also significantly reduced in squirrels treated with cortisol. Despite these profound shifts in the functionality of the HPA axis, squirrel body mass, offspring survival, and adult survival were unaffected by experimental increases in cortisol concentrations. Our results highlight that even short-term experimental increases in glucocorticoids can affect adrenal gland functioning and CBG concentrations but without other side-effects.

## Introduction

Associations between glucocorticoids (GCs) and life history or behavioral traits are increasingly studied, due to their role as a mechanistic link between the genome and the environment, and to uncover general relationships between hormones and fitness (Breuner et al., 2008; Dantzer et al., 2016). GCs are released by the hypothalamic-pituitary-adrenal (HPA) axis in response to environmental challenges and have widespread effects on physiology and behavior (Sapolsky et al., 2000; Romero, 2004). Correlational studies have helped advance our understanding of the relationships between GCs and phenotypic traits in wild animals, but establishing the causality of such relationships requires experimental manipulation of GCs. In laboratory settings, hormone manipulations are logistically feasible (e.g. Karatsoreos et al., 2010; Lussier et al., 2009), but experimental studies conducted in wild populations are likely to provide better insights into the ecologically relevant effects of GCs on life history variation.

Hormone manipulations in wild animals are more challenging than in the laboratory, but several methods have been developed (see Sopinka et al., 2015). That said, exogenous GCs may have unintended physiological side effects, which may influence or skew interpretation of the results obtained from manipulative studies. One potential problem with hormone manipulations is related to the fact that the endocrine system is a homeostatic system that is controlled by negative feedback mechanisms and tends to compensate for disruption. Therefore, if animals are treated with a hormone, the endogenous production of the hormone may be reduced after a few days, and longer treatment duration may lead to the regression of the endocrine gland, and have important consequences for endocrine homeostasis (Fusani, 2008). Such effects are well documented in humans, as both cortisol and synthetic GCs (which may be more potent, see Meikle and Tyler, 1977), are used to treat a range of ailments (Arabi et al., 2010; Kirwan et al., 2007). For example, cortisol administration may lead to side-effects, including suppression of the HPA axis and reduced adrenal function (Broide et al., 1995; Feiwel et al., 1969; Jacobs et al., 1983). Although such side-effects are usually temporary (Morris and Jorgensen, 1971; Streck and Lockwood, 1979), in extreme cases, patients may develop more severe and long term conditions such as Cushing’s syndrome (see Axelrod, 1976) or secondary adrenal insufficiency that can lead to Addison’s disease (Arlt and Allolio, 2003).

If GC manipulations affect the adrenal glands, endogenous production of GCs, and endocrine homeostasis, this may lead to unintended consequences in wild animals. This could jeopardize the value of performing such studies, as they could adversely influence survival and reproduction. Indeed, some studies indicate that elevations in GCs reduce estimates of fitness (Bonier et al., 2009; Breuner et al., 2008; Wingfield et al., 1998), but it is unclear if this is due to an unintended complication from the manipulation rather than a natural consequence of increased GCs. Although these issues were highlighted 10-15 years ago (Romero, 2004; Fusani, 2008; see also Sopinka et al. 2015; Crossin et al., 2016), detailed studies about the potential complications of manipulating hormones in wild animals have not been widely performed except in birds. Torres-Medina et al. (2018) reviewed the consequences of experimentally elevated GCs on baseline and stress-induced corticosterone levels from previous studies on multiple bird species that were published 2005-2015. Many but not all of these studies were conducted in free-living birds. They showed that most studies that experimentally elevated GCs (using silastic implants, time release pellets, or osmotic pumps) examined how corticosterone treatment affected baseline corticosterone levels but very few investigated treatment effects on stress-induced corticosterone levels. Their results documented that birds treated with exogenous GCs exhibited lower stress-induced corticosterone levels, suggesting that experimental elevation of GCs can suppress the activity of the HPA axis in wild birds just as in studies in humans or laboratory rodents.

Unlike birds, studies that experimentally elevated GCs in free-living mammals are extremely rare and we are not aware of any study that has investigated both how treatment with GCs affects the HPA axis in wild mammals or how changes in the HPA axis induced by treatment with exogenous GCs affects fitness proxies. We examined how exogenous cortisol affected the HPA axis and life history traits of North American red squirrels (*Tamiasciurius hudsonicus*) by treating squirrels with exogenous cortisol or control vehicle. We expected that exogenous cortisol would increase fecal glucocorticoid metabolite (FGM) and plasma cortisol concentrations, but that, as may be the case in humans receiving GC therapy or birds with corticosterone implants, HPA axis responsiveness would decrease. We examined how administration of exogenous cortisol affected the responsiveness of the HPA axis by measuring the change in plasma cortisol concentrations following 1) capture and handling and 2) pharmaceutical suppression with dexamethasone (Dex) and pharmaceutical stimulation with adrenocorticotropin hormone (ACTH; hereafter “Dex/ACTH challenges”). Because we expected that administration of cortisol would suppress HPA axis responsiveness, we also examined how quickly the HPA axis recovered after administration of exogenous cortisol by measuring HPA axis responsiveness (using the Dex/ACTH challenges) in squirrels that received exogenous cortisol on the same day of sampling or the day after their last treatment. We expected that exogenous cortisol might lead to increased body mass in squirrels (Axelrod, 1976), but did not expect our treatment dosages to be sufficiently high to cause anorexia through sustained adrenal impairment (Arlt and Allolio, 2003). As we aimed to keep GCs within a physiologically-relevant (‘normal’) range for this species, we did not expect to see negative effects of our treatments on body mass or adult or litter survival.

## Methods

### Study population

All of our research was approved by the *Animal Care and Use Committee* at the University of Michigan (PRO00005866). We studied a natural population of red squirrels in the Yukon, Canada (61°N, 138°W) that has been monitored since 1987 (Boutin et al., 2006; McAdam et al., 2007). All squirrels in this population are individually identified by a unique ear tag in each ear, as well as a unique color combination of colored wires attached to each ear tag which allow researchers to identify individuals from a distance. Squirrels were live-trapped (Tomahawk Live Trap Co., WI, USA), during which they were weighed using a Pesola spring balance, and fecal samples were collected from underneath traps, placed on ice, and stored at −20 °C upon return to the field station (Dantzer et al., 2010). Female and male reproductive condition was assessed through palpation or by expressing milk from the teats in females (see McAdam et al., 2007).

### Experimental manipulations of GCs

In 2015 and 2016, squirrels were randomly allocated to either control (8 g all natural peanut butter, 2 g wheat germ, no cortisol) or cortisol treatments (8 g peanut butter, 2 g wheat germ with 6, 8, or 12 mg of cortisol [H4001, Sigma Aldrich, USA]). Dosages of 0, 6, 8, and 12 mg cortisol were selected following previous studies in red squirrels (Dantzer et al., 2013) and laboratory rodents (Casolinia et al., 1997; Catalani et al., 2002; Mateo, 2008) that used similar dosages to induce a moderate increase in GCs. Treatments were provided directly to squirrels by putting the treatment in a bucket that was hung from trees on their territories (Dantzer et al., 2013). To ensure that target squirrels (identifiable through ear tags/radio-collars) were consuming the treatments, camera traps (Reconyx PC900 HyperFire Professional Covert IR) were placed by the buckets of 31 squirrels for five continuous days. Out of 155 d of camera trapping, conspecific pilferage was only observed ten times, and there was one case of heterospecific pilferage by a grey jay (*Perisoreus canadensis*). Consumption of each treatment was estimated daily by checking buckets for any leftovers and estimating these as a percentage. Squirrels consumed on average 91.9% of their total peanut butter treatments (median = 100%, SD = 12.13%, range = 43-100%).

### Effects of exogenous cortisol on fecal glucocorticoid metabolites (FGM)

To evaluate the effects of cortisol treatments on FGM, fecal samples were collected in 2015 and 2016 from male and female squirrels fed with 0, 6, 8, and 12 mg cortisol/day (Table 1). Glucocorticoid metabolites from fecal samples were extracted and assayed as previously validated and described (Dantzer et al., 2011, 2010) using a 5α-pregnane-3β,11β,21-triol-20-one enzyme immunoassay (Touma et al., 2003). Intra- and inter-assay CVs for pools diluted 1:250 (n = 13 plates) were 7.4% and 15.4%. For pools diluted 1:500 (n = 13 plates) this was 7.5% and 17.9%. Pools diluted 1:100 (n = 9 plates) had intra and inter-assay CVs of 10.0% and 17.9%, and for pools diluted at 1:700 (n = 9 plates) this was 6.4% and 18.9%. Samples from control (n = 135) or cortisol (n = 237) treated squirrels included those collected before (range: 0-21 days before treatment started, mean ± SE: 7 ± 0.7 days), during, and after treatment (range:1-21 days after treatment, mean ± SE: 9.4 ± 0.8 days, Table 1).

### Effects of exogenous cortisol on plasma cortisol concentrations and corticosteroid binding capacity

Non-breeding male squirrels were fed cortisol (8 mg/day) or control treatments for one (n = 40 squirrels) or two weeks (n = 26). The time that squirrels consumed their treatments was estimated by checking buckets at regular intervals between 25 minutes and a few hours (shown as hours:min, mean = 1:53 hrs, SD = 1:12 hrs). Squirrels were either blood sampled the same day they consumed their last treatment (n = 36 squirrels, mean = 3:30 hrs after treatment consumption, range = 0:57-8:55 hrs) or the day after they consumed their last treatment (n = 30, mean = 22:46 hrs after treatment consumption, range = 14:57-30:55 hrs). Note that for 19 next day bleed squirrels, the time of treatment consumption was not recorded. Blood samples obtained within 3 min of squirrels entering a trap (n = 54 samples) are referred to as *true baseline* samples (Romero et al. 2005). If the first blood sample was obtained >3 min after squirrels went into traps (n = 12), this is referred to as *stress-induced* samples. Stress-induced samples are considered to reflect the effects of capture and handling on plasma cortisol concentrations or were taken to see if dexamethasone administration (see below) would reduce plasma cortisol concentrations. Although the time of day we obtained blood samples varied, there was no daytime sampling bias between cortisol treated (between 9:41 and 18:07, mean = 13:23) and control squirrels (between 9:33 and 17:31, mean = 13:50, t-test, t_42.15_ = 1.14, p = 0.26). We also conducted a general linear model containing time of day of sampling (b = −0.02, SE = 0.18, t_14_ = −0.12, P = 0.91) and a quadratic term for time of day (b = −0.41, SE = 0.24, t_14_ = −1.68, P = 0.11) and found no significant effect of sampling time on plasma cortisol.

Blood samples were obtained from the nailbed and collected into heparinized capillary tubes, and plasma was separated via centrifugation and frozen at −20° C. Total plasma cortisol concentrations were assayed using an ImmuChem coated tube cortisol radioimmunoassay (MP Biomedicals, New York, USA) following the manufacturer’s instructions, with the exception that, due to small sample volumes, plasma and tracer volumes of 12.5 µL and 500 µL were used. This assay has already been validated and used to measure plasma cortisol in other rodent species (Karatsoreos et al. 2010; Brooks and Mateo, 2013). We validated this kit by first showing linearity and then demonstrating that the assay reliably responds to changes in HPA axis activity through the decreases in plasma cortisol we observed in response to Dex and the increases in plasma cortisol in response to ACTH (Fig. 3). Linearity was tested by pooling several samples and serially diluting these from 1 (neat) to 1:64. Results were plotted, visually inspected, and evaluated with linear regression (*R*^2^ adj = 0.991, p < 0.001). According to the manufacturer, the assay detection limit is 1.7 ng/ml, and samples that read below this value (n = 8) were set at 1.7 ng/ml. Most samples were run in duplicate, but because of small plasma volumes only one estimate was obtained for 33.9% of samples. Average standard and sample intra-assay CVs were 7.9% (n = 4 assays). Inter-assay CVs for the five standards provided (10, 30, 100, 300 and 1000 ng/ml cortisol) were 11.1%, 15.4%, 8.8%, 4.0% and 7.7%.

Corticosteroid binding capacity (CBG) was measured in plasma stripped of endogenous steroids using dextran-coated charcoal (DCC) and diluted to a final dilution of 1/50 in phosphate buffered saline with 0.1% gelatin (PBS). Three tubes (final volume of 150 µL) were prepared for each sample: two containing 160 nM cortisol (10% 1,2,6,7-^3^H-cortisol, Perkin Elmer, Waltham, MA, and 90% non-labeled cortisol, C-106, Sigma-Aldrich) to measure total binding, and one containing an additional 4 µM non-labeled cortisol to measure nonspecific binding (primarily by albumin). After incubating tubes overnight, 300 µL of ice-cold DCC was added and left for 15 minutes to strip free cortisol from the plasma mixture. The tubes were then centrifuged at 2000 × *g* at 4 °C for 12 minutes. The supernatant (containing bound cortisol) was decanted into scintillation vials, to which 4 mL of scintillation fluid (Emulsifier-Safe cocktail, Cat. No. 6013389, Perkin Elmer, Groningen, Netherlands) was added. Vials were counted in a scintillation counter. Specific binding by CBG was calculated by subtracting nonspecific binding counts from total binding counts. Specific binding scintillation counts were converted to nM binding by measuring the total counts in the 150 µL of the 160 nM solution and adjusting for the plasma dilution. Some CBG-bound hormone is lost to the DCC during the 15 minute DCC exposure. Using pooled plasma exposed to DCC for 5-20 minutes, we calculated the rate of loss of CBG-bound cortisol (data not shown). From this, we calculated that the 15 minute DCC exposure resulted in the loss of 28.5% of CBG-bound hormone, and all our specific binding measurements were corrected accordingly. To calculate the percent free cortisol, we estimated free cortisol concentrations (i.e. not bound by CBG) using the total cortisol concentration, the equilibrium dissociation constant for red squirrels of 61.1 nM (Delehanty et al., 2015), and the equation in Barsano and Baumann (1989). As plasma volumes were limited, only 58 samples could be assayed for both CBG/percent free cortisol and total cortisol (see below for details).

### Effects of exogenous cortisol on HPA axis reactivity

To determine how our cortisol treatments affected the responsiveness of the HPA axis, we used two different methods. First, we assessed the response to capture and handling (“handling stress”) in cortisol-treated and control squirrels by acquiring a series of blood samples starting immediately after they entered the live-trap. Squirrels were bled at intervals of 0-3, 3-6, 6-12 and 18-22 min after trap doors closed either the same day as confirming they ate their last treatment (mean time elapsed = 3:36 hrs, range = 0:57-8:55 hrs, cortisol n = 10, control n = 13) or the next day (mean time elapsed = 22:46, hrs range = 14:57-30:55 hrs, cortisol n = 5, control n = 6).

Second, in a separate set of squirrels, we assessed how cortisol-treated and control squirrels (n = 32) responded to intra-muscular injections of dexamethasone (Dex, a glucocorticoid receptor antagonist) and adrenocorticotropic hormone (i.e., Dex/ACTH challenges). We used previously described protocols (Boonstra and McColl, 2000), with modified concentrations of Dex (3.2 mg/kg) and ACTH (4 IU/kg). Briefly, squirrels were captured and a true baseline blood sample (0-3 min after entering trap) was obtained followed by the acquisition of a second blood sample (stress-induced sample) an average of 12:37 min (min:sec, n = 13, range = 5:00-28:00 min) after the squirrel entered the trap (time of the stress-induced sample was not recorded for 13 different squirrels as traps were checked ∼60 min after they were set). Squirrels were then injected with Dex and a third blood sample was acquired 60 min after injection. Following the acquisition of the third blood sample, squirrels were injected with ACTH and then blood sampled 30 and 60 min after ACTH injection.

### Effects of exogenous cortisol on adult squirrel body mass

We assessed how treatment with exogenous cortisol affected body mass of non-breeding females (n = 21 females, 64 body mass measures before treatment and 85 records during treatment) and non-breeding males (n = 47 males, 37 body mass measures before treatment and 28 records during treatment) by live-trapping them approximately once per week and weighing them to the nearest 5g with a spring scale (McAdam et al., 2007). We compared non-breeding squirrel body mass in cortisol treated and control squirrels sampled in 2015 and 2016 before and during the treatments.

### Effects of exogenous cortisol on litter survival

As a part of our long-term data collection, we track the reproduction of females during pregnancy and lactation by capturing them, palpating their abdomens to identify pregnancy stage, and expressing milk from their teats to identify if they are lactating (McAdam et al., 2007). We also retrieve pups from their natal nest soon after parturition (“first nest entry”) and approximately 25 d after parturition (“second nest entry”) to collect a range of data described elsewhere (McAdam et al., 2007). We used these data to identify how treatment with exogenous cortisol affected litter survival in females treated only during pregnancy (n = 71) and in a separate group of females treated only during lactation (n = 17) compared to controls. In these comparisons, we also included data on litter fate collected in 2012 from squirrels fed the same dosages (0, 6, 12 mg cortisol/day) for similar periods of time (see Dantzer et al., 2013). When females gave birth, their nests were located an average of 1.6 d after parturition (SD = 1.6 d, range = 0-6 days) and then again 25.5 d after parturition (SD = 1.5 d, range = 21-29 d).

We examined how our treatments affected whether females treated during pregnancy (control n = 24, 6 mg cortisol/day n = 9, 8 mg cortisol/day n = 16, 12 mg cortisol/day n = 22) or lactation (control n = 8, 12 mg cortisol/day n = 9) lost their litters before the first nest entry (via abdominal palpation) or between the first and second nest entry (indicated by cessation of lactation). Females treated during pregnancy were treated from the estimated last third of pregnancy (based on abdominal palpation), until five days post parturition (treatment duration range = 8–25 days, mean = 18, SD = 4). Females treated during lactation were treated for 10 continuous days, from days 5 to 15 post parturition (mean = 9.9 d, SD = 0.6, range = 8 −11 days).

### Effects of exogenous cortisol on adult squirrel survival

The effects of treatment with exogenous cortisol on survival of adult squirrels was monitored through regular live trapping and behavioral observations (McAdam et al., 2007). Survival data were only available from squirrels studied in 2015 (n = 50, including 41 females and nine males). These squirrels were fed either control treatments (n = 25, 10-26 days, mean = 19, SD = 7) or 12 mg cortisol/day (n = 25, 8-35 days, mean = 20, SD = 7). Although we did not know the ages of all squirrels, there was no age bias between squirrels fed control (eight known ages, mean = 4.05, SD = 1.05 years) and those fed cortisol (nine known ages, mean = 3.97, SD = 0.87 years, t_13.8_ = 0.18, p = 0.86). We estimated survival until exactly 1 year after the treatments were stopped.

### Statistical analyses

Analyses were conducted using R statistical software (v 3.3.3, R Core Team, 2017). When there were multiple measures for individual squirrels, linear mixed-effects models (LMMs) were conducted using packages ‘lme4’ (v 1.1.10, Bates et al., 2015) and all such models contained ‘squirrel ID’ as a random intercept term. If there were no repeated measures, general linear models (GLM) were used. To make comparisons between groups, we used the ‘glht’ function in R package ‘multcomp’ (Hothorn et al., 2017). Model residuals were plotted to check for conformity with homogeneity of variance and normality (Zuur et al., 2010). Where necessary, data were ln transformed. Regression lines were visualized using R package ‘visreg’ (v 2.2.2, Breheny and Burchett, 2016).

We tested effects of treatments on FGM concentrations using LMMs, analyzing female and male data separately due to differences in reproductive states. Models for females included dose (0, 6, 8, 12 mg of cortisol/day), reproductive state (non-breeding, pregnant, lactating), Julian date, and whether the squirrel was treated on the sampling day (yes/no) as fixed effects, with an interaction term for dose and treatment (yes/no). Models for males included the same variables (but only doses of 0 and 8 mg) except reproductive state (all were non-breeding).

To assess how our treatments affected the responsiveness of the HPA axis, we used two separate LMMs to assess if cortisol treated and control squirrels differed in their plasma cortisol concentrations following 1) our capture and handling stress experiments where we obtained a series of blood samples 2 to 28 min after the trap doors closed and 2) our Dex/ACTH challenges. For the LMM to assess the effects of capture and handling on plasma cortisol concentrations, the model included a fixed-effect for treatment (control or cortisol) and the time taken to acquire the blood sample expressed in minutes since the squirrel was trapped (standardized following Schielzeth, 2010). For the LMM to assess plasma cortisol concentrations following the Dex/ACTH challenges, the model included the fixed effect (control or cortisol) and a categorical variable for when the blood sample was obtained (‘true baseline’, ‘stress-induced, ‘60 minutes after Dex injection’ [hereafter; DEX], ‘30 minutes after ACTH injection’ [hereafter; ACTH30], and ‘60 minutes after ACTH injection’ [hereafter; ACTH60]). Two plasma samples with very low binding (<10%) were excluded from the analysis. Some models included squirrels treated for either 1 or 2 weeks, and some included squirrels that were treated in both spring and autumn (n = 18 squirrels, with treatments switched between periods, with the exception of two squirrels fed GCs twice and one squirrel fed control treatments twice). Where this was the case, treatment duration (1/2 weeks) and whether or not squirrels had been treated before (yes/no) were included in our initial models. Because these two variables (treated for one/two weeks, and whether or not squirrels had been treated previously) were not significant in any of the models, we do not discuss them below.

To assess effects of treatments on CBG concentrations and percent free cortisol, we subset samples collected at different intervals after squirrels entered the traps (effects of handling stress samples) and those from Dex/ACTH challenges. For our handling stress samples, data from samples collected on the same day (n = 12) and the day after (n = 3) the last treatment was consumed were pooled. This model included an interaction between the sampling day (same/next) and treatment (control or 8 mg cortisol/day). Due to limited data (only 58 samples were analyzed for CBG, across all categories), only the effects of treatment (control or 8 mg cortisol/day) on CBG and percent free cortisol were tested for squirrels Dex/ACTH challenged on the same day as consuming their last treatments (n = 16 squirrels). Models for squirrels ACTH challenged the day after consuming their last treatments (n = 27 squirrels) included interactions between sample time (stress-induced, Dex, ACTH30, ACTH60) and treatment (control or 8 mg cortisol/day).

To estimate the total plasma cortisol in a 24 h period, true baseline cortisol was plotted against the time since treatment was consumed. Regression line equations were used to calculate the area under these lines for both control and cortisol treated squirrels, using the ‘trapzfun’ command in package ‘pracma’ (Borchers, 2018), and areas under the curve were compared with χ^2^ tests.

Data on body mass were subset into those collected in spring (non-breeding females fed 0 or 12 mg cortisol/day) and autumn (non-breeding males fed 0 or 8 mg cortisol/day). Body masses were compared using LMMs including a two-way interaction between treatment, and time (before/during treatment). To assess differences between litter survival (lost/not lost), and adult survival (yes/no) GLMs were applied using binomial errors. Models included a binary fixed effect for treatment (12 mg cortisol/day or control) and sex (only for adult survival). Because the duration of the treatments varied among different squirrels, we also included total days of treatment and an interaction between treatment and treatment duration in all these models to assess how our treatments affected body mass and adult or litter survival. Dispersion parameters (using R package blemco, Korner-Nievergelt et al., 2015) between 0.75 and 1.4 were taken to accept overdispersion was not problematic.

## Results

### Effects of treatments on fecal glucocorticoid metabolite concentrations

Overall, squirrels fed cortisol treatments (6, 8, 12 mg/day) had significantly higher FGM concentrations than when they were not being fed, but the magnitude of increase depended on the dosage (F_3,263.8_ = 11.5, p < 0.001, Fig. 1). Both female and male control squirrels fed plain peanut butter had similar FGM concentrations when they were being fed their treatments compared to when they were not being fed their treatments (females: b = −0.03, SE = 0.15, z = 0.22, p = 1.0; males: b = 0.14, SE = 0.36, z = 0.41, p = 0.96). FGM concentrations in both females and males fed 6, 8, or 12 mg cortisol/day were significantly higher compared when they were being fed compared to when they were not being fed (6 mg: b = 0.78, SE = 0.28, z = 2.8, p = 0.032; 8 mg: females: b = 0.79, SE = 0.21, z = 3.9, p < 0.001; males: b = 1.37, SE = 0.49, z = 2.82, p = 0.013; 12 mg: b = 1.39, SE = 0.17, z = 8.0, p < 0.001). Concentrations of FGM during treatment in female squirrels treated with 12 mg vs 8 mg (b = 0.39, SE = 0.28, z = 1.4, p = 0.63), 6 mg vs 8 mg (b = 0.36, SE = 0.36, z = 1.0, p = 0.88), and 6 mg vs 12 mg cortisol/day (b = 0.74, SE = 0.35, z = 2.1, p = 0.18) were not significantly different. Julian date did not affect FGM concentrations in females (F_1,284.9_ = 3.44, p = 0.06) or males (F_1,15.2_ = 0.73, p = 0.41). Reproductive condition did not affect FGM in this dataset, possibly because of limited sample numbers on some reproductive states (see Table 1, F_2,105.0_ = 1.37, p = 0.26).

**Fig 1.**
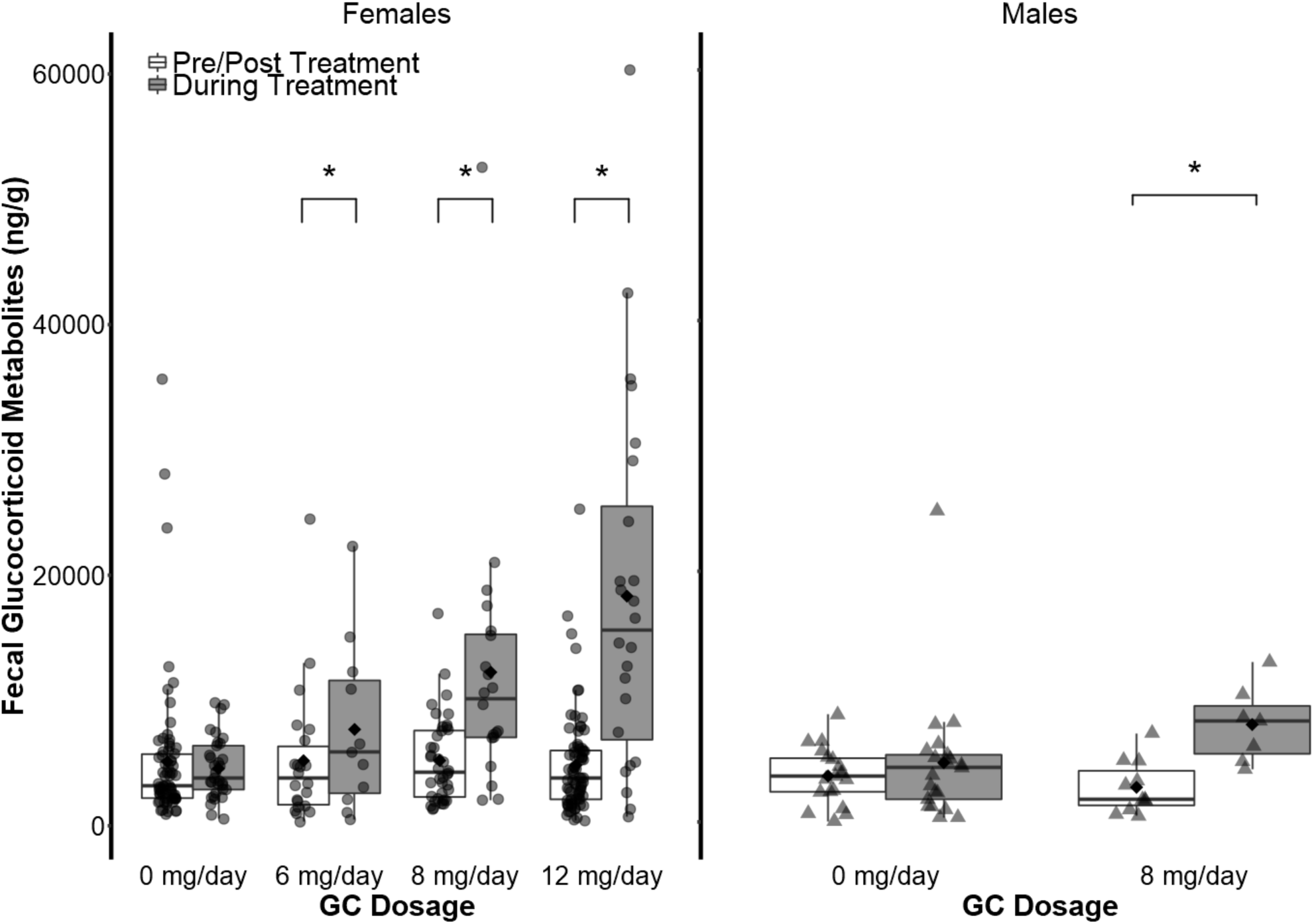
FGM concentrations in squirrels treated with control (0 mg/day) or with cortisol (6, 8, or 12 mg/day) treatments. Asterisks (*) indicate significant differences (p < 0.05). Upper and lower hinges correspond to the first and third quartiles. Upper/lower whiskers extend from the hinge to the highest/lowest value that is within 1.5x the interquartile range. White diamonds indicate means.

### Effects of treatments on total plasma cortisol concentrations over 24 hr period

We plotted true baseline plasma cortisol concentrations against the time since treatment was consumed to estimate the area under these lines for both control and cortisol treated squirrels. The area under these lines was used to estimate the total plasma cortisol concentrations over a 24 hr period. Overall, we estimated that cortisol treated squirrels experienced significantly higher plasma cortisol (total area = 7907.4 units) than controls (total area = 4110.8 units) over a 24 h period (χ^2^ = 614.4, DF = 1, p < 0.001, Fig. 2).

**Fig. 2.**
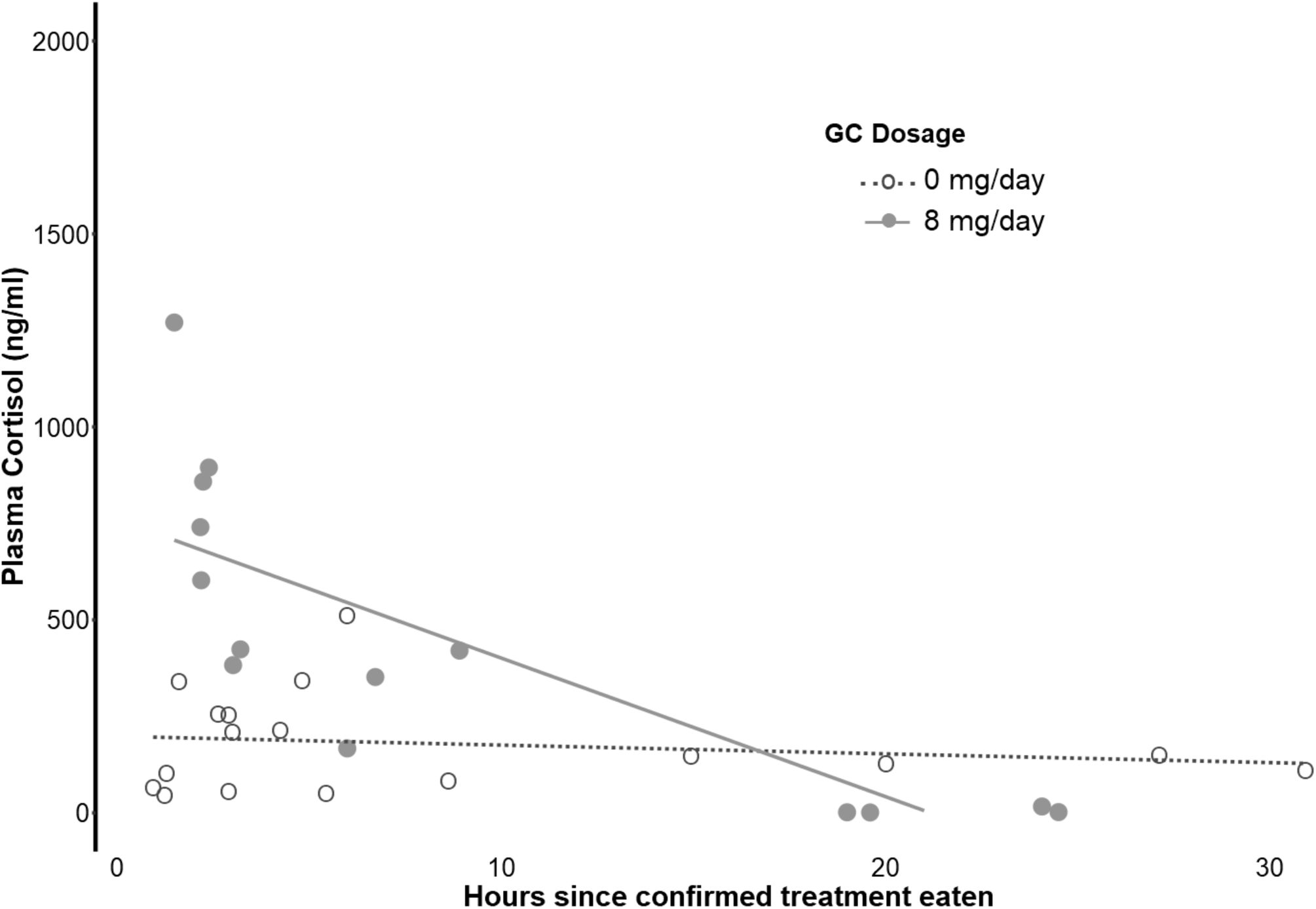
True baseline plasma cortisol concentrations in squirrels fed cortisol (GC, 8 mg/day) and controls (0 mg/day) sampled between 1 to 31 hours after confirming they consumed their last treatments. Different points correspond to different individual squirrels.

### Responsiveness of HPA axis to capture and handling in squirrels sampled same day or day after last treatment

The effects of capture and handling on plasma cortisol concentrations were significantly different between control and cortisol treated squirrels, in addition to whether the squirrels were sampled on the same day or day after their last treatment. In squirrels sampled the same day as receiving their last treatment, plasma cortisol concentrations were generally higher in cortisol treated squirrels compared to controls, but their responsiveness to capture and handling differed (Fig. 3A). In control squirrels sampled the same day as receiving their last treatment, plasma cortisol concentrations significantly increased as handling time increased (b = 0.31, SE = 0.11, t = 2.8, DF = 61.4, p = 0.007, Fig. 3A) whereas they declined as handling time increased in cortisol treated squirrels (b = −0.56, SE = 0.17, t = − 3.3, DF = 61.4 p = 0.002, Fig. 3A).

**Fig. 3.**
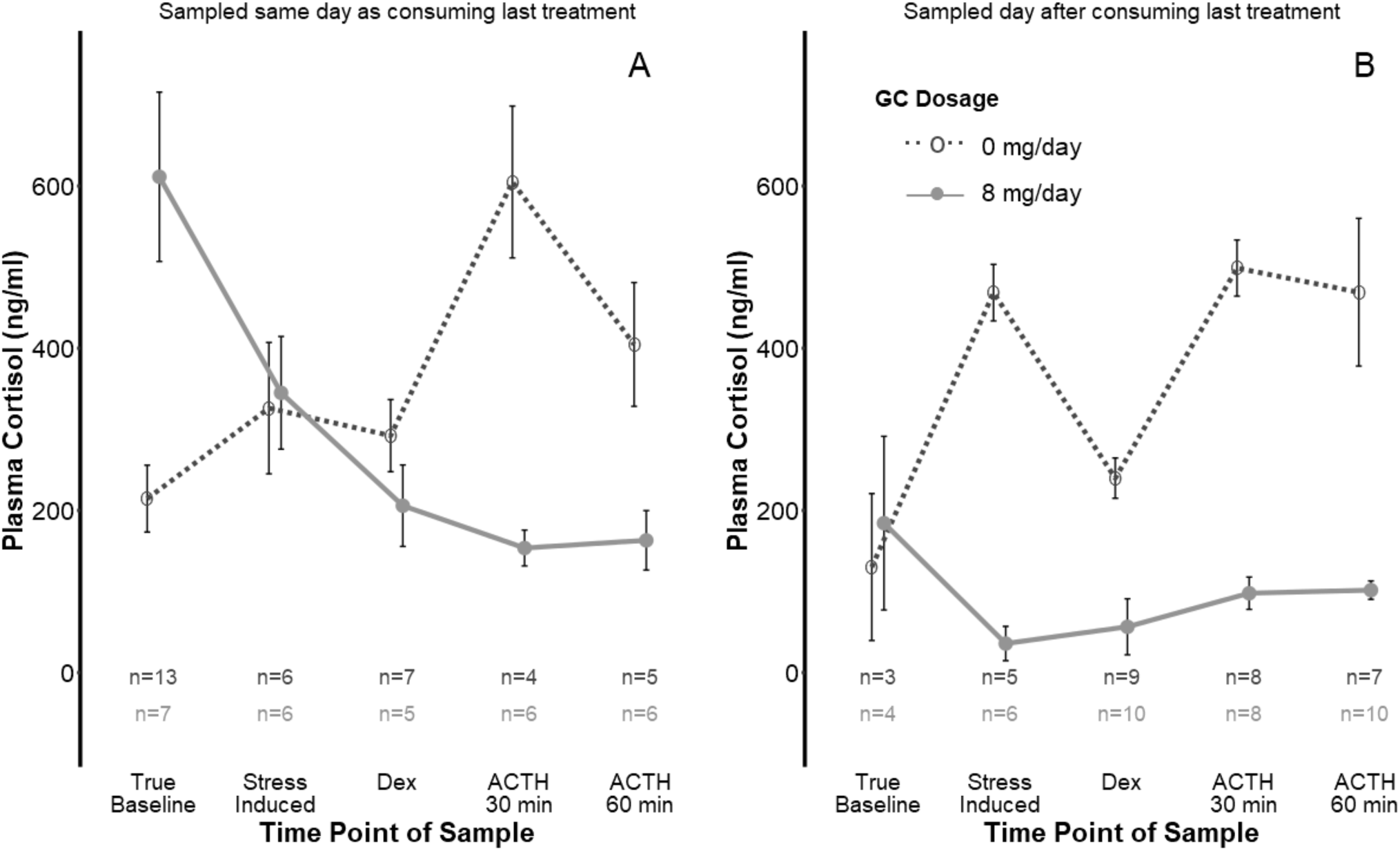
Effect of time elapsed since squirrels entered traps (“handling stress”) on plasma cortisol concentrations from control squirrels (0 mg/day), or those treated with GCs (8 mg/day) and (A) sampled the same day as consuming their last treatment or (B) sampled the day after consuming their last treatment. Mean and SE are shown.

**Fig 4.**
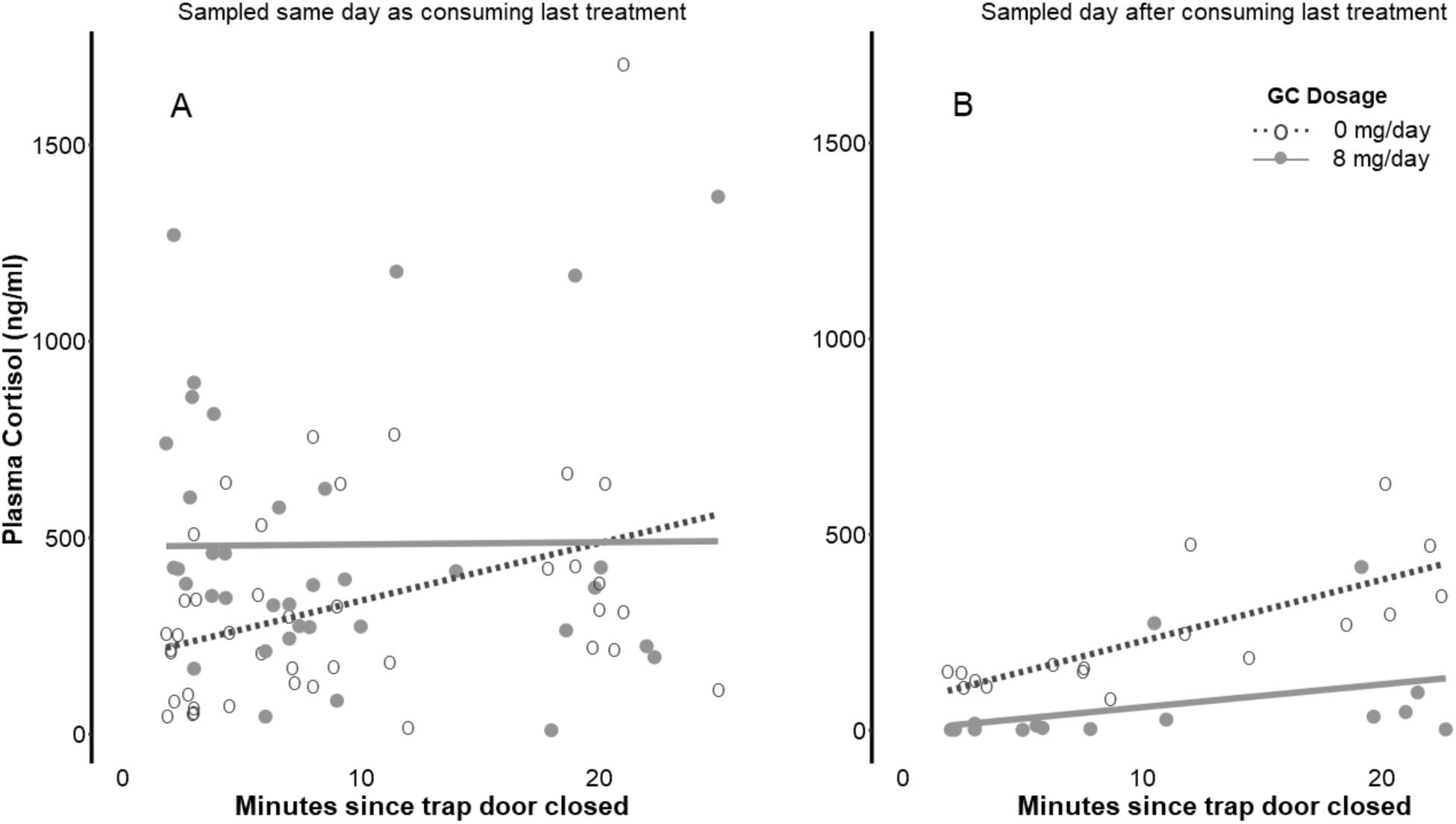
Plasma cortisol concentrations following Dex/ACTH challenges conducted on male squirrels treated with 0 mg (Control) or 8 mg cortisol/day for 7-14 days (GCs). A) Males were trapped the same day as confirming they consumed their last treatment. B) Males were trapped the day after being fed their last treatment, but note that the time they consumed their treatments was not recorded. Note that true baseline samples for cortisol treated squirrels were highly variable (two samples of 321.2 and 412.7 ng/ml, and two of 1.7 ng/ml). Means and standard errors are shown.

**Fig. 5.**
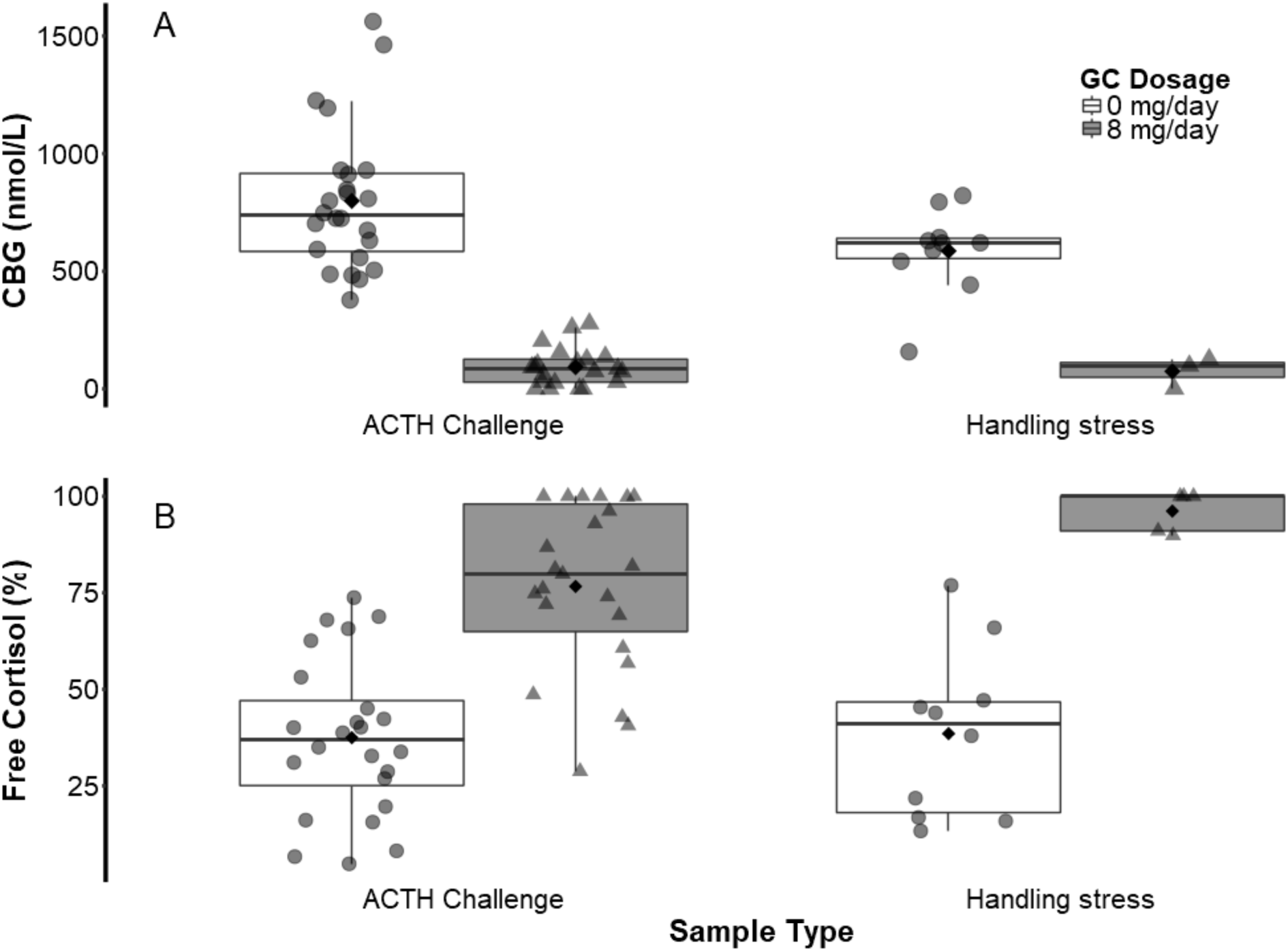
A) Plasma corticosteroid binding globulin (CBG) and B) percent free cortisol in control and cortisol treated squirrels subjected to both Dex/ACTH challenges (Dex/ACTH Challenge) and response to handling stress (blood samples obtained 0-3, 3-6, 6-12 and 18-22 minutes after entering trap were lumped together). The figure includes squirrels that were sampled both on the same day as consuming their last treatment and the day after consuming their last treatment. Upper and lower hinges correspond to the first and third quartiles. Upper/lower whiskers extend from the hinge to the highest/lowest value that is within 1.5x the interquartile range. White diamonds indicate means.

In squirrels sampled the day after receiving their last treatment, plasma cortisol concentrations were generally lower in cortisol treated squirrels than in control squirrels (b = −1.8, SE = 0.81, t = −2.2, DF = 8.4, p = 0.057, Fig. 3B), though this difference was not significant. Handling time increased plasma cortisol concentrations in both control and cortisol treated squirrels (b = 0.42, SE = 0.20, t = 2.1, DF = 19.2, p = 0.048) and this was not affected by treatment (b = 0.40, SE = 0.1.4, DF = 18.7, p = 0.16, Fig. 3B).

### Responsiveness of HPA axis to Dex/ACTH challenges in squirrels sampled on the same day as consuming last treatment

In squirrels sampled on the same day as consuming their last treatment, HPA axis responsiveness to our Dex/ACTH challenges differed between control and cortisol treated squirrels (F_9,35.5_ = 6.4, p < 0.001, Fig. 3A). Squirrels treated with cortisol (8 mg/day) had significantly higher true baseline cortisol concentrations (611.4 ± 104.4 ng/ml) than control squirrels (214.5 ± 41.3 ng/ml, b = 394.7, SE = 88.7, z = 4.4, p < 0.001, Fig. 3A) but cortisol treated and control squirrels had similar stress-induced plasma cortisol concentrations (b = 15.9, SE = 114.5, z = 0.14, p = 1, Fig. 3A). Both cortisol treated (205.8 ± 50.2 ng/ml) and control (292.2 ± 44.5 ng/ml) squirrels responded to Dex, as indicated by the reductions in their plasma cortisol concentrations 60 min after the Dex injection compared to stress-induced plasma cortisol concentrations, although these reductions in were not significant (control: b = −33.9, SE = 81.2, z = − 0.44, p = 1.0; cortisol treated: b = −139.6, SE = 88.7, z = −1.57, p = 0.62). Control squirrels had significantly higher plasma cortisol concentrations in samples taken 30 minutes after ACTH injection compared to those obtained 60 min after the Dex injection (604.9 ± 93.6 ng/ml, b = 321.1, SE = 81.2, z = 3.97, p < 0.001) but not in samples taken 60 min after ACTH injection (404.6 ± 76.3 ng/ml, b = −213.4, SE = 85.1, z = −2.51, p = 0.10). In cortisol treated squirrels, plasma cortisol concentrations were unaffected by ACTH as plasma cortisol concentrations in samples taken 30 minutes after ACTH injection (153.5 ± 22.1 ng/ml, b = −51.8, SE = 88.7, z = −0.58, p = 1.0) and 60 minutes after ACTH injection (163.1 ± 36.7 ng/ml, b = 9.5, SE = 83.5, z = 0.11, p = 1.0) were no different from those obtained 60 min after the Dex injection.

### Responsiveness of HPA axis to Dex/ACTH challenges in squirrels sampled the day after consuming their last treatment

In squirrels sampled the day after consuming their last treatment, HPA axis responsiveness to our Dex/ACTH challenges differed between control and cortisol treated squirrels (F_4,46.5_ = 9.2, p < 0.001, Fig. 3B). Cortisol treated squirrels that were sampled the day after consuming their last treatment had lower plasma cortisol concentrations than controls at all sampling times (Fig. 3B). Although true baseline plasma cortisol concentrations did not differ between cortisol treated and control squirrels (b = 48, SE = 82.7, z = 0.6, p = 0.99, Fig. 3B), stress-induced plasma cortisol concentrations (35.9 ± 21.2 ng/ml) were on average 92.3% lower in cortisol treated squirrels than in control squirrels (468.5 ± 34.8 ng/ml, b = −404.8, SE = 67.4, z = −6.0, p < 0.001, Fig. 3B). Plasma cortisol concentrations after the Dex injection were an average of 76.4% lower in cortisol treated squirrels (56.5 ± 34.6 ng/ml) than controls (239.8 ± 24.8 ng/ml, b = −183.3, SE = 53.7, z = −3.4, p = 0.006). Cortisol treated squirrels had plasma cortisol concentrations (mean = 97.9 ± 19.9 ng/ml) that were on average 80.4% lower than in control squirrels (mean = 498.8 ± 34.6 ng/ml, b = 398.1, SE = 57.5, z = −6.9, p < 0.001) 30 min after the ACTH injection. This difference remained 60 min after the ACTH injection (cortisol mean = 101.7 ± 11.4 ng/ml, control mean = 469.0 ± 89.2 ng/ml, a difference of 78.3%, b = −368.2, SE = 56.8, z = −6.5, p < 0.001, Fig. 3B).

In squirrels sampled the day after receiving their last treatment, control squirrels (b = −211.9, SE = 53.7, z = −3.9, p < 0.001), but not cortisol treated squirrels (b = 9.6, SE = 49.4, z = 0.2, p = 1.0), had significantly lower plasma cortisol concentrations 60 minutes after Dex injections compared to stress-induced concentrations. Thirty minutes after the ACTH injection (ACTH30), control squirrels (b = 257.3, SE = 45.6, z = 5.6, p < 0.001), had significantly higher plasma cortisol concentrations than 60 minutes after Dex injections. In cortisol treated squirrels, plasma cortisol concentrations were higher at 30 minutes after ACTH injection (ACTH30) than 60 minutes after Dex injection, but this was not significant (b = 42.6, SE = 44.7, z = 0.95, p = 0.95, Fig. 3B).

### Effects of cortisol treatment on plasma CBG and free cortisol concentrations

Squirrels treated with cortisol had significantly lower CBG concentrations in plasma samples obtained during our Dex/ACTH challenges compared to controls regardless of whether the samples were obtained on the same day as consuming their last treatments (F_1,5.0_ = 51.0, p < 0.001) or the day after consuming their last treatment (F_1,12.9_ = 29.3, p < 0.001, Fig. 6A). Consequently, squirrels treated with cortisol had significantly higher free cortisol concentrations in plasma samples obtained during the Dex/ACTH challenges regardless of whether they were sampled on the same day (F_1,5.0_ = 7.6, p = 0.04) or the day after consuming their last treatments (F_1,12.1_ = 15.3, p = 0.002, Fig. 6B). CBG concentrations in plasma samples obtained during the Dex/ACTH challenges from squirrels sampled the day after consuming their last treatment did not vary among the different sample types (stress-induced, Dex, ACTH30, ACTH60: F_3,6.2_ = 1.0, p = 0.46), nor did percent free cortisol (F_3,12.3_ = 0.5, p = 0.71).

In plasma samples obtained from squirrels at regular intervals after they entered our live traps, CBG concentrations were significantly lower in plasma samples acquired from cortisol treated squirrels compared to controls (F_1,7.0_ = 24.5, p = 0.002, Fig. 6A), but percent free cortisol did not differ between cortisol treated and control squirrels (F_7.0_ = 2.3, p = 0.17). The effect of treatment on plasma CBG concentrations (F_7.0_ = 1.3, p = 0.30) and percent free cortisol (F_1,7.0_ = 1.9, p = 0.21) was not different between squirrels sampled the same day or the day after last treatment.

### Effects of treatments on body mass

Both control and cortisol treated females (effect of treatment period, b = 11.9, SE = 3.7, t87.4 = 3.26, p = 0.001) but not males (b = 1.33, SE = 3.2, t32.4 = 0.41, p = 0.68) were heavier when they were being treated (n = 23 females, n = 11 males) compared to before they were being treated (n = 24 females, n = 17 males). However, control and cortisol treated females and males did not differ in body mass while they were being treated, as indicated by the lack of significant interactions between treatment and treatment period in both females (b = −3.83, SE = 4.9, t_87.2_ = −0.79, p = 0.43) and males (b = 6.5, SE = 4.84, t_29.5_ = 1.34, p = 0.19). Although females and males in both treatment groups (control or cortisol) were treated for varying lengths of time, treatment duration did not affect body mass as indicated by the lack of significant interactions between treatment and treatment duration for both females (b = −1.36, SE = 12.1, t_17.8_ = −0.11, p = 0.91) and males (b = 1.73, SE = 3.8, t_49.7_ = 0.45, p = 0.65).

### Effects of treatments on litter survival

For females that were treated during pregnancy, there was no significant difference in litter survival rates before the first nest entry between females treated with cortisol (6, 8, 12 mg/day) during pregnancy (14/47 litters lost) and controls (7/24 litters lost; z = 0.28, p = 0.78). Similarly, there were also no significant difference in litter survival between the first and second nest entry for females treated with cortisol during pregnancy (17/47 litters lost) and controls (4/24 litters lost: z = 0.01, p = 0.99). Even though control (range = 11-23 d) and cortisol treated (range = 8-25 d) pregnant females were treated for different lengths of time, there was no effect of treatment duration for both control and cortisol treated females on litter survival prior to the first nest entry (treatment × treatment duration, z = −0.39, p = 0.69) and between the first and second nest entry (z = −0.16, p = 0.87).

For females treated during lactation (n = 17), there was no significant difference in litter survival between the first and second nest entry, with 1/8 control females losing their litter and 3/9 cortisol fed (12 mg/day) females losing their litter (z = 0.001, p = 1). Although treatment duration varied for control (range = 10-11 d) and cortisol treated (range = 8-10 d) lactating females, there was once again no effect of treatment duration for both control and cortisol treated females (treatment × treatment duration, z = −0.005, p = 0.97).

### Effects of treatments and treatment duration on adult survival

There was no difference in survival to one year following cessation of the treatments between controls (18/25 survived to one year) and those fed cortisol (14/25 survived to one year; z = −1.09, p = 0.27). Although the duration of the cortisol or control treatments slightly varied among cortisol treated (range = 8-35 d) or control (range = 10-26 d) squirrels, treatment duration did not affect adult survival as indicated by the lack of significant interaction between treatment and treatment duration (z = 0.63, p = 0.53). There was no difference in survival between males (7/9 survived to one year), and females (25/41 survived to one year, z = 1.19, p = 0.24).

## Discussion

Our results on cortisol manipulations in wild red squirrels, spanning a range of dosages, life history stages, and including both sexes, provide important information regarding the response of wild animals to such hormone manipulation including the possible fitness consequences of hormone manipulation. Squirrels treated with cortisol had higher FGM concentrations and true baseline plasma cortisol concentrations over a 24 h period. However, similar to studies in humans, laboratory rodents, or wild birds (see Introduction), our results highlight that exogenous GCs can cause the adrenals to stop responding to handling stress or pharmaceutical (Dex/ACTH) challenges. Although we documented that treatment with exogenous GCs affected the responsiveness of the HPA axis, these effects were short-lived and did not affect fitness proxies, including body mass, and offspring or adult survival.

Our results indicate that concentrations of CBG were significantly reduced in squirrels treated with cortisol for one or two weeks, suggesting that chronically elevated GCs reduce CBG concentrations. This reduction in CBG may result in a higher bioavailability of plasma cortisol (Breuner et al., 2013). Previous studies in rats have shown that administration of exogenous GCs can inhibit the rate of CBG production and secretion in the liver (Feldman et al., 1979), and one study found that, 24 hours after acute stress, CBG concentrations were reduced in rats (Fleshner et al., 1995). Chronic elevations in GCs has also been shown to lead to reduced CBG concentrations in most species studied to date (Armario et al., 1994; Breuner et al., 2013). A previous study found that CBG concentrations in red squirrel plasma started to decrease as quickly as four hours after the start of Dex/ACTH challenges, suggesting that although high concentrations of CBG may buffer squirrels from the effects of high concentrations of free cortisol caused by acute stressors, these concentrations decline rapidly when the duration of the stressor is longer than a few hours (Boonstra and McColl, 2000). However, this does not seem to carry any noticeable cost as there were no changes in body mass or litter and adult survival in response to our treatments.

Previous reviews have emphasized the importance of maintaining hormone concentrations within a physiological range when performing hormone manipulations (Crossin et al., 2016; Fusani, 2008; Zera, 2007). Studies in mammals have shown that acute experimental challenges can cause increases in plasma cortisol that are comparable to those achieved by our treatments. For example, in both laboratory rats and wild animals, physical restraint, open field trials, and maze tests may cause >10 fold increases in plasma GCs (Cockrem, 2013). In our study, the highest recorded true baseline plasma cortisol concentration in cortisol treated squirrels was approximately seven times higher than the average control true baseline plasma cortisol concentration. This indicates that the increase in plasma cortisol caused by our 8 mg cortisol/day treatment is within the physiological range for a squirrel. However, it is possible that the duration of elevated plasma cortisol caused by our treatments is longer than that caused by a natural ecological factor that elicits an increase in plasma cortisol. Studies in rats show that plasma GCs increase quickly in response to acute stress, but returns to baseline concentrations within 2-5 hours after the stressor is removed (Marin et al., 2007; Mizoguchi et al., 2001), but in cortisol treated squirrels, plasma cortisol remained elevated, compared to control squirrels, for an estimated 17 hours post-treatment (Fig. 2).

Our results highlight the importance of regularly provisioning individuals with treatments to sustain increases in hormone concentrations. Although we found that squirrels fed cortisol had significantly higher concentrations of plasma cortisol over a 24 hr period than the controls, it was important to provision individuals with the treatments every 24 hrs. This was because plasma cortisol in cortisol treated squirrels did decrease to concentrations well below those of control squirrels >20 hrs after consuming their treatments. When it is feasible, daily supplementation may be effective in maintaining sustained elevations in hormone concentrations and provide an alternative to other methods like implants that carry some disadvantages (Sopinka et al., 2015). For example, Torres-Medina et al. (2018) showed that silastic implants, time release pellets, or osmotic pumps that contain corticosterone can also suppress the responsiveness of the HPA axis in birds, which could decrease overall exposure to circulating corticosterone. Our study shows that even regular provisioning of exogenous GCs rather than implants, such as through food in our study (see also Dantzer et al., 2017) or through other methods (Vitousek et al. 2018), may also decrease the activity of the HPA axis and cause a reduced ability to mount an increase in circulating GCs in response to an environmental challenge.

Our results also highlight the potentially adverse consequences that may occur when ending hormone manipulations in wild animals. When sampled less than a day after the end of cortisol treatment, squirrels did not respond to handling stress or ACTH injection, and appeared to have impaired adrenal function and lower CBG concentrations. Data collected from cortisol treated squirrels the day after consuming their last treatment showed that plasma cortisol was very low compared to control squirrels, suggesting exogenous GCs have been excreted but endogenous GCs were being produced at a lower rate than in control animals. However, plasma cortisol did increase with handling stress, suggesting some recovery of adrenal function within 24 hrs of stopping the treatments. Our results suggest that the adrenal gland may need time to recover from treatment, and endogenous cortisol production may not return to pre-treatment levels for several days.

Hormone manipulations can provide powerful tools to study relationships between hormones and life history traits, and in recent years methods have been developed to achieve this (Sopinka et al., 2015). Many studies aim to experimentally elevate GCs to test the “cort-fitness hypothesis”, which proposes that elevations in baseline GCs decreases survival or reproduction (Bonier et al., 2009), or to document the effects of elevated GCs on behavior or life history traits (Crossin et al., 2016). We show that elevation of plasma cortisol concentrations within the physiological range for 1-2 weeks had profound effects on measures of HPA axis reactivity and CBG concentrations. This is similar to the results of Torres-Medina et al. (2018) that showed that treatment with corticosterone implants can also cause reduced corticosterone levels in response to capture and handling. However, our study took the result from Torres-Medina et al. (2018) one step further as we showed how treatment with exogenous GCs suppresses CBG concentrations and we also investigated the possibility of fitness consequences of reduced HPA axis reactivity. Despite these observed shifts in the functionality of the neuroendocrine stress axis and the sustained elevations in GCs, we found no change in body mass or offspring and adult survival. This indicates that some species can tolerate bouts of increased GCs and rapid reorganization of the stress axis without negatively impacting survival and reproduction.

